# Predicting Peptide-MHC Binding Affinities with Imputed Training Data

**DOI:** 10.1101/054775

**Authors:** Alex Rubinsteyn, Timothy O'Donnell, Nandita Damaraju, Jeff Hammerbacher

## Abstract

Predicting the binding affinity between MHC proteins and their peptide ligands is a key problem in computational immunology. State of the art performance is currently achieved by the allele-specific predictor NetMHC and the pan-allele predictor NetMHCpan, both of which are ensembles of shallow neural networks. We explore an intermediate between allele-specific and pan-allele prediction: training allele-specific predictors with synthetic samples generated by imputation of the peptide-MHC affinity matrix. We find that the imputation strategy is useful on alleles with very little training data. We have implemented our predictor as an open-source software package called MHCflurry and show that MHCflurry achieves competitive performance to NetMHC and NetMHCpan.

## I. Introduction

In most vertebrates, cytotoxic T-cells enforce multi-cellular order by killing infected or cancerous cells. Each organism possesses a poly-clonal army of T-cells which collectively are able to distinguish unhealthy cells from healthy ones. This amazing feat is achieved through the winnowing and expansion of clonal T-cell populations possessing highly specific T-cell receptors (TCRs) (1). Each distinct TCR recognizes a small number of similar peptides bound to an MHC molecule on the surface of a cell (2). Though there are many steps in “antigen processing” (3), it has become apparent that MHC binding is the most restrictive step. Peptide-MHC affinity prediction is the well-studied problem of predicting the binding strength of a given peptide and MHC pair (4). Early approaches focused on “sequence motifs”(5), followed by regularized linear models, linear models with interaction terms such as SMM with pairwise features (6), and more recently the NetMHC family of predictors, a collection of related models based on ensembles of neural networks. Two of these predictors, NetMHC (7) and NetMHCpan (8), have emerged as the methods of choice across multiple fields of study within immunology, including virology (9), tumor immunology (10), and autoimmunity (11).

NetMHC is an *allele-specific* method which trains a separate predictor for each allele's binding dataset, whereas NetMHCpan is a *pan-allele* method whose inputs are vector encodings of both a peptide and a subsequence of a particular MHC molecule. The conventional wisdom is that NetMHC performs better on alleles with many assayed ligands whereas NetMHCpan is superior for less well-characterized alleles (12).

In this paper we explore the space between *allele-specific* and *pan-allele* prediction by imputing the unobserved values of peptide-MHC affinities for which we have no measurements and using these imputed values for pre-training of allele-specific binding predictors.

## 2. Data and evaluation metrics

Two datasets were used from a recent paper studying the relationship between training data and pMHC predictor accuracy(13). The training dataset (BD2009) contained entries from IEDB (14) up to 2009 and the test dataset (BLIND) contained IEDB entries from between 2010 and 2013 which did not overlap with BD2009 (Table 1).

**Table I.** Train (BD2009) and test (BLIND) dataset sizes.

|  | Alleles | IC50 Measurements | Expanded 9mers |
| --- | --- | --- | --- |
| BD2009 | 106 | 137,654 | 470,170 |
| BLIND | 53 | 27,680 | 83,752 |

Throughout this paper we will evaluate a pMHC binding predictor using three different metrics:

- **AUC**:Area under the ROC curve. Estimates the probability that a “strong binder” peptide (affinity≤500nM) will be given a stronger predicted affinity than one whose ground truth affinity is > 500nM.
- **F_1_score**:Measures trade-off between sensitivity and specificity for predicting “strong binders” with affinities≤500nM.
- **Kendall's***τ*: Rank correlation across the full spectrum of binding affinities.

## 3. Comparison of imputation algorithms as predictors

A dataset of peptide-MHC affinities for *n* peptides and *a* alleles may be thought of as a n × a matrix where peptide/allele pairs without measurements are missing values. The task of predicting values at these positions is known as matrix completion or imputation (depending on the community and data source). We investigated the performance of several imputation algorithms as a standalone solution to the peptide-MHC affinity prediction problem. The algorithms considered were:

- **meanFill**:Replace each missing pMHC binding affinity with the mean affinity for that allele. This is a very simple imputation method which serves as a baseline against which other methods can be compared.
- **knnlmpute** (15):Each missing entry *X_ij_* is imputed using the values in the *k* closest columns with observation in row *i*. Similarity between alleles is computed as 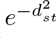, where *d_st_* is the mean squared difference between observed entries of alleles *s* and *t*.
- **svdlmpute** (15):Imputation using iterative fixed rank SVD decomposition.
- **softlmpute** (16): A singular value thresholding method which iteratively estimates a low-rank matrix completion without forcing the pre-specification of a particular solution rank. Instead, the *softlmpute* method is parameterized by a shrinkage value λ that is subtracted from each singular value.
- **MICE** (17):Average multiple imputations generated using Gibbs sampling from the joint distribution of columns.

We evaluated the performance of these methods using three-fold cross validation on BD2009, only considering peptides which occurred in at least three alleles and excluding alleles with less than five measurements (Table 2). All imputation methods were implemented in the *fancyimpute* Python library (18). Since MICE outperformed the other methods on two of the three predictor metrics, we selected it for the subsequent neural network experiments.

**Table 2.** Cross-validation performance of imputation algorithms on BD2009 dataset

| Imputation Method | Parameter | | AUC | $F_1$ score | Kendall's $\tau$ |
| --- | --- | --- | --- | --- | --- |
| meanFill |  |  | 0.67665 | 0.04950 | 0.17675 |
| knnImpute | $k = 1$ | | 0.80907 | 0.57952 | 0.40201 |
| | $k = 3$ | | 0.83189 | 0.57594 | 0.42086 |
| | $k = 5$ | | 0.83103 | 0.56118 | 0.41703 |
| MICE | $n = 25$ | | 0.85861 | 0.57597 | 0.44978 |
| | $n = 50$ | | **0.86127** | 0.56527 | **0.45944** |
| softImpute | $\lambda = 5$ | | 0.78981 | 0.39158 | 0.33408 |
| | $\lambda = 10$ | | 0.83248 | 0.53575 | 0.39763 |
| | $\lambda = 20$ | | 0.85608 | **0.60599** | 0.43754 |
| svdImpute | rank = 5 |  | 0.82305 | 0.57040 | 0.39117 |
|  | rank = 10 |  | 0.83667 | 0.58433 | 0.40048 |
|  | rank = 20 |  | 0.82986 | 0.57038 | 0.38817 |

## 4. Neural network architecture

Each MHCflurry predictor is a feed-forward neural network containing (1) an embedding layer which transforms amino acids to learned vector representations, (2) a single hidden layer with *tanh* nonlinearity, (3) a sigmoidal scalar output. This network is implemented using Keras (19).

Three-fold cross validation on the training set was used to select the hyper-parameters. The best model had 32 output dimensions for the amino acid vector embedding, a hidden layer size of 64, a dropout rate of 50%, and 250 training epochs. These hyper-parameters achieved reasonable performance across alleles, but it's likely that performance could be further improved by setting the hyper-parameters separately for each allele.

## 5. Data encoding

Like the NetMHC family of predictors (20), MHCflurry uses fixed length 9mer inputs which requires peptides of other lengths to be transformed into multiple 9mers. Shorter peptides are mapped onto 9mer query peptides by introducing a sentinel “X” at every possible position, whereas longer peptides are cut into multiple 9mers by removing consecutive stretches of residues. The predicted affinity for a non-9mer peptide is the geometric mean of the predictions for the generated 9mers. When *n* training samples derive from a single non-9mer sequence then their weights are adjusted to 1/n.

We map IC50 concentrations onto a regression targets between 0.0 and 1.0 using the same scheme as NetMHC, *y*=1.0 − *max* (1.0, log_50000_(*IC*50)).

## 6. Training

For each allele, we train a MHCflurry model using the measured peptide affinities for the allele and the values imputed by MICE based on other alleles in the training set. As training progresses, we place quadratically decreasing weight on the imputed values.

A randomly generated peptide is unlikely to bind a given MHC strongly, but a data acquisition bias toward strong binders in the training set can lead models to assign a high affinity to most peptides. As a form of regularization, we augment the training set at each epoch to include random peptides with affinity set to be maximally weak. The number of random negative peptides is 20% of the training size (without imputation). At each training epoch, a fresh set of random peptides is generated.

## 7. Results

We evaluated the effect of imputation by drawing subsets of the BD2009 training set for the well-characterized allele HLA-A*02:01. Predictors were trained on a range of simulated training set sizes and tested on the remaining data (Figure 1). We find that imputation gives a modest improvement up to approximately 100 training samples. With more training data there is no benefit to imputation.

**Fig1.**
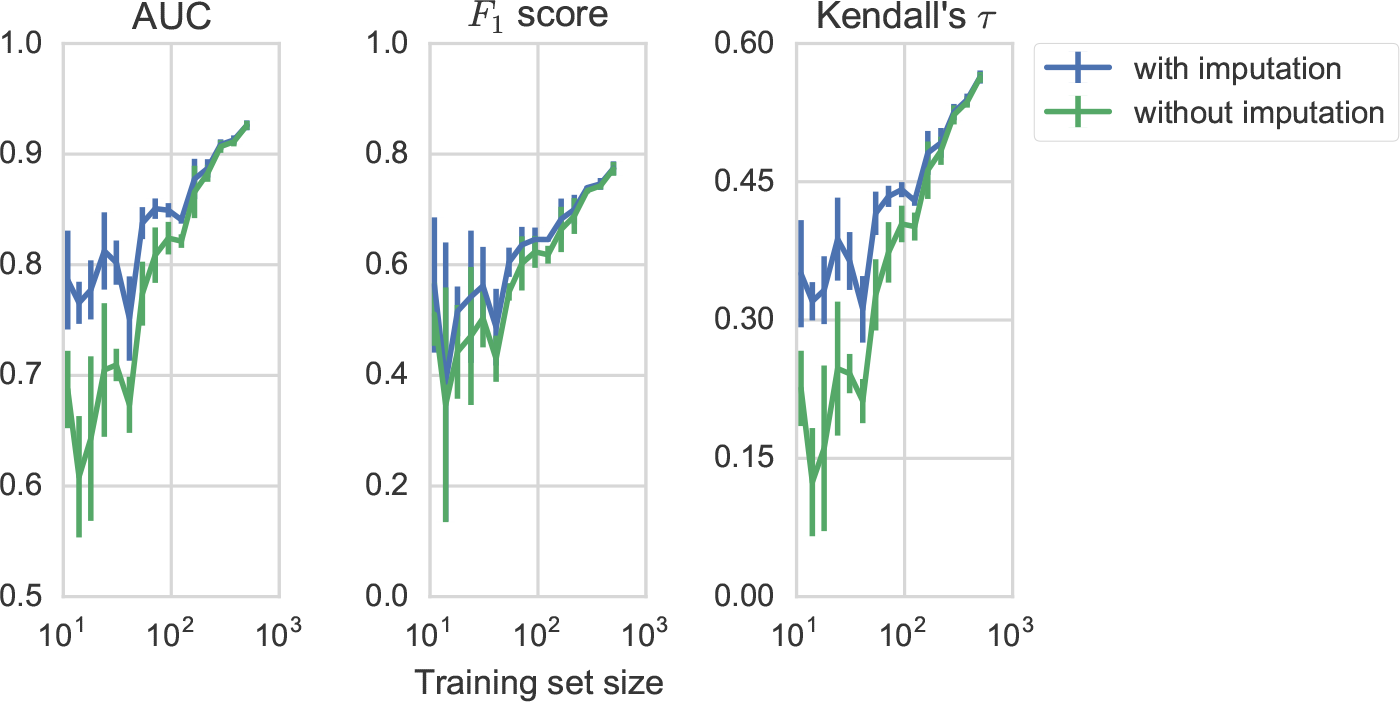
MHCflurry performance on down-sampled training data for HLA-A*02:01 with and without imputation

We then compared the performance of MHCflurry against NetMHC, NetMHCpan, and SMM on the blind test data. The MHCflurry ensemble model contains 32 predictors initialized with different random weights. The MHCflurry ensemble is competitive with NetMHC and NetMHCpan.

**Table 3.**
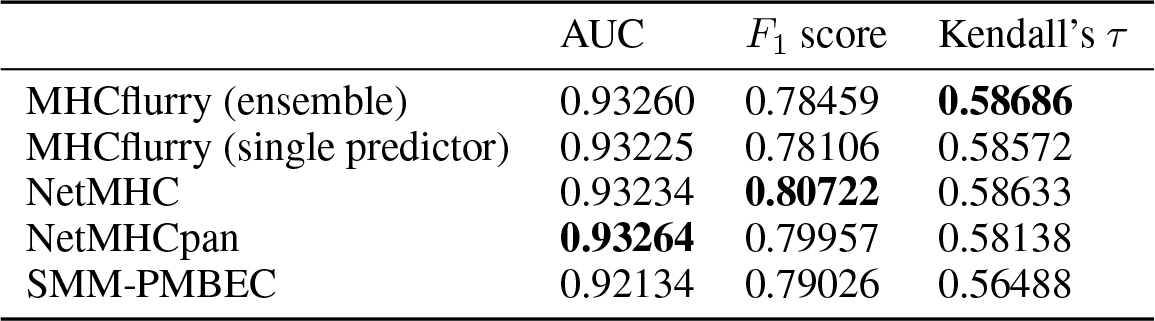
Performance on BLIND dataset

## 8. Discussion

Imputing training data shows promise in cross-validation as a way to improve performance on alleles with few observations, but only seems to help for very small training sizes (≤ 100). Unfortunately, none of the alleles included in the BLIND dataset had fewer than 100 samples in BD2009, and only one had fewer than 200. Thus, additional work is required to assess the accuracy of MHCflurry and other predictors on alleles with scarce training data. Additionally, we need to further investigate the interaction between imputation parameters, the decay schedule for the weights of imputed samples, and stopping criteria for training individual allele-specific predictors.

## 9. Code

MHCflurry is available at https://github.com/hammerlab/mhcflurry The data, scripts, and notebooks used to generate the plots and tables in this paper are available at https://github.com/hammerlab/mhcflurry-icml-compbio-2016/.

